# Lip movements enhance speech representations and effective connectivity in speech dorsal stream and its relationship with neurite architecture

**DOI:** 10.1101/2021.09.27.462075

**Authors:** Lei Zhang, Yi Du

## Abstract

Lip movements facilitate speech comprehension, especially under adverse listening conditions, but the neural mechanisms of this perceptual benefit at the phonemic and feature levels remain unclear. This fMRI study addresses this question by quantifying regional multivariate representation and network organization underlying audiovisual speech-in-noise perception. We found that valid lip movements enhanced neural representations of phoneme, place of articulation, or voicing feature of speech differentially in dorsal stream regions, including frontal speech motor areas and supramarginal gyrus. Such local changes were accompanied by strengthened dorsal stream effective connectivity. Moreover, the neurite orientation dispersion of left arcuate fasciculus, a structural basis of speech dorsal stream, predicted the visual enhancements of neural representations and effective connectivity. Our findings provide novel insight to speech science that lip movements promote both local phonemic and feature encoding and network connectivity in speech dorsal pathway and the functional enhancement is mediated by the microstructural architecture of the circuit.

## Introduction

Speech perception becomes challenging in noisy environments and older adults (Du, Buchsbaum, Grady, & Alain, 2016; L. Zhang, Fu, Luo, Xing, & Du, 2021). However, in face-to-face communication, we routinely extract visual speech cues from the speaker’s articulatory movements, which substantially benefits speech comprehension, especially in challenging listening conditions (Ross, Saint-Amour, Leavitt, Javitt, & Foxe, 2007; Sumby & Pollack, 1954) and in hearing impaired senior populations (Puschmann et al., 2019). This multisensory perceptual gain has been validated across speech hierarchies from isolated syllables to continuous speech, but it may operate in distinct ways (Grant & Seitz, 1998).Visual speech relays correlated information about “when” the speaker is saying (the timing of the acoustic signal, influencing attention and perceptual sensitivity) and supplementary information about “what” the speaker is saying (place and manner of articulation, constraining lexical selection) (Peelle & Sommers, 2015). For continuous speech, the temporal coherence between the area of mouth opening and speech envelope facilitates the attentive tracking of the speaker, signals temporal markers to segment words or syllables, or provides linguistic cues, thereby improving speech intelligibility (Grant & Seitz, 1998; Hauswald, Lithari, Collignon, Leonardelli, & Weisz, 2018; Park, Kayser, Thut, & Gross, 2016). For speech syllables and words, visual lip movements provide the place and manner of articulation to constrain lexical competition (Grant & Walden, 1996). The visual speech head start processed before speech vocalization is thought to increase the precision of articulatory prediction (Karas et al., 2019). Growing magnetoencephalography (MEG) and electroencephalogram (EEG) studies have emphasized on neural entrainment and encoding of continuous speech under the audiovisual context (Crosse, Butler, & Lalor, 2015; Crosse, Di Liberto, & Lalor, 2016; Giordano et al., 2017; Keitel, Gross, & Kayser, 2020; Park, Ince, Schyns, Thut, & Gross, 2018), that largely advances our understanding of multisensory speech processing. However, direct observation of where in the brain and how valid visual speech information modulates the focal neural representations of phonemes (the most fundamental linguistic unit) and articulatory-phonetic features, as well as the network organization during speech-in-noise perception, is still lacking. Moreover, it remains unknown which neuroanatomical structure undergirds functional changes underlying the visual enhancement of speech-in-noise perception.

Previous MEG and EEG research on continuous speech has shown that visual speech improves the neural tracking of speech envelope (Crosse et al., 2015; Giordano et al., 2017), and facilitates the neural encoding of both spectrotemporal and phonetic features of speech (O’Sullivan, Crosse, Di Liberto, de Cheveigné, & Lalor, 2021). The visual benefit on neural tracking of speech was stronger under noisy conditions than quiet conditions, demonstrating the inverse effectiveness in audiovisual speech processing (Crosse et al., 2016). Despite the limitation of spatial resolution, recent MEG studies started to locate the brain regions involved in the visual enhancement of speech encoding, including the left motor cortex and inferior frontal gyrus (IFG) (Giordano et al., 2017; Keitel et al., 2020). The left posterior superior temporal gyrus/sulcus (pSTG/S) has been implicated as another critical region in audiovisual speech integration in functional magnetic resonance imaging (fMRI) studies (Erickson, Heeg, Rauschecker, & Turkeltaub, 2014; Nath & Beauchamp, 2011) and intracranial EEG studies (Karas et al., 2019; Micheli et al., 2020). However, the left pSTG/S is recently found to represent the common redundant features of the bimodal signals, whereas left speech motor areas represent the synergistic feature of them (Park et al., 2018). Moreover, the neural entrainment to lip movements in the left motor cortex (Park et al., 2016) and enhanced effective connectivity between frontal motor and temporal cortices (Giordano et al., 2017) were correlated with the visual benefit on speech comprehension. Those findings are consistent with the model suggesting that the speech dorsal stream, including the left pSTG/S, supramarginal gyrus (SMG), and speech motor areas (IFG and premotor/motor cortex), is involved in integrating visual and auditory speech in addition to auditory and visual cortices (Bernstein & Liebenthal, 2014). Considering that frontal speech motor areas are engaged to a higher extent in adverse listening conditions to provide articulatory predictions to compensate for degraded bottom-up speech processing (Alain, Du, Bernstein, Barten, & Banai, 2018; Du, Buchsbaum, Grady, & Alain, 2014; Du et al., 2016; Nuttall, Kennedy-Higgins, Hogan, Devlin, & Adank, 2016; Pickering & Garrod, 2013; Skipper, Devlin, & Lametti, 2017), we hypothesized that visual lip movements would promote functional activities mainly along the dorsal stream when speech is degraded by noise. Also, we are interested in whether visual speech cues would shape neural representations of phonemes and articulatory-phonetic features differentially in distinct regions and how the network connectivity would be changed accordingly.

Here, we adopted the fMRI technique to specify the univariate and multivariate neural activities and effective connectivity patterns when subjects discriminated audiovisual consonant-vowel syllables under different signal-to-noise ratios (SNRs) with and without valid lip movements. Behaviorally, valid visual cues significantly improved phoneme identification via facilitating the recognition of place of articulation but not voicing regardless of SNR. Univariate analysis showed that right auditory and motor areas and bilateral visual regions were more activated when subjects were viewing valid visual cues. Multivariate pattern analysis (MVPA) revealed better neural representations of speech phonemes with valid visual cues in left speech motor areas and SMG. Interestingly, those regions exhibited distinct representational improvements by visual speech cues that the classification of voicing was enhanced in the left opercular part of IFG (IFG_op_) while the classification of place of articulation was improved in the left inferior part of precentral gyrus (PrCG_inf_) and SMG. This is the first evidence that lip movements sharpened neural encoding of phonemes by predicting and constraining selective articulatory features in distinct dorsal stream regions. Next, we carried out the dynamic causal modeling (DCM) analysis to investigate the influence of lip movements on network organization involved in audiovisual speech perception. Bidirectional connectivity between SMG/AG (angular gyrus) and frontal speech motor areas (Broca’s area and PrCG_inf_) and top-down connectivity from SMG/AG to sensory areas (auditory and visual cortices) were enhanced, while bottom-up connectivity from auditory cortex to SMG/AG was inhibited with valid visual cues. These results suggest the auditory dorsal stream as a crucial pathway in audiovisual speech integration, which led us to further exam the relationship between the white matter basis of the speech dorsal stream, i.e., the arcuate fasciculus (AF) (Friederici, 2017; Hickok & Poeppel, 2007), and functional changes of visual enhancement. We used state-of-art neurite orientation dispersion and density imaging (NODDI) technique, which constructs a three-compartment tissue model with the multi-shell high angular resolution diffusion-weighted imaging (HARDI) data (Zhang, Schneider, Wheeler-Kingshott, & Alexander, 2012), to quantify the fine-grained microstructural neurite morphology of the left AF. We found that a greater visual enhancement of phoneme representations in the left IFG_op_ and a stronger visual enhancement of top-down connectivity from speech motor areas (Broca’s area and PrCG_inf_) to the auditory cortex were correlated with a higher neurite orientation dispersion of the left AF. Our findings provide novel evidence of both local phonemic and feature representations and network connectivity changes underlying the visual enhancement of speech-in-noise perception, and for the first time link individual microstructural variations of structure connectivity with functional activity during audiovisual speech processing.

## Results

Participants (N = 24) were presented with 4 consonant-vowel syllables (/ba/, /da/, /pa/, /ta/) organized into 2 orthogonal articulatory features, place of articulation (bilabial: /ba/ and /pa/; lingua-dental: /da/ and /ta/) and voicing (voiced: /ba/ and /da/; voiceless: /pa/ and /ta/). Syllables were embedded into a speech spectrum-shaped noise at −8, 0, and 8 dB SNRs and paired with matching lip movements videos or still closed mouth pictures in visual valid (VV) and visual invalid (VI) conditions, respectively (see Materials and Methods). We measured whole-brain activity using fMRI while subjects listened to and identified the audiovisual syllables. HARDI data were recorded in addition to task fMRI.

### Lip movements improved recognition of place of articulation at the behavioral level

We replicated previous findings (Grant & Walden, 1996) that visual speech provides place of articulation but not voicing to improve speech-in-noise identification. As shown in Fig. 1A, the main effects of visual validity and SNR on phoneme identification accuracy were both significant (visual validity: *F*(1, 23) = 391.72, *P* < 0.001, 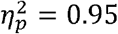; SNR: *F*(2, 46) = 31.43, *P* < 0.001, 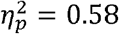, repeated-measures analysis of variance (ANOVA)) without a significant interaction (*F*(2, 46) = 0.42, *P* = 0.658, 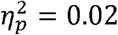). However, valid visual cues did not promote the recognition of voicing (e.g., if the stimulus /ba/ was identified as /da/, it was scored correct for voicing) (*F*(1, 23) = 0.00, *P* = 0.976, 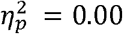, Fig. 1C), which confirms that the perception of voicing was determined by the auditory modality. In contrast, as shown in Fig. 1B, valid visual cues significantly improved the recognition of place of articulation (e.g., if the stimulus /ba/ was identified as /pa/, it was scored correct for place) (*F*(1, 23) = 433.83, *P* < 0.001, 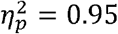), and the SNR effect on the identification of place was insignificant under the VV condition (*F*(2, 23) = 0.13, *P* = 0.879, 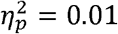). This confirms that the recognition of place was determined mainly by the visual modality.

**Fig. 1.**
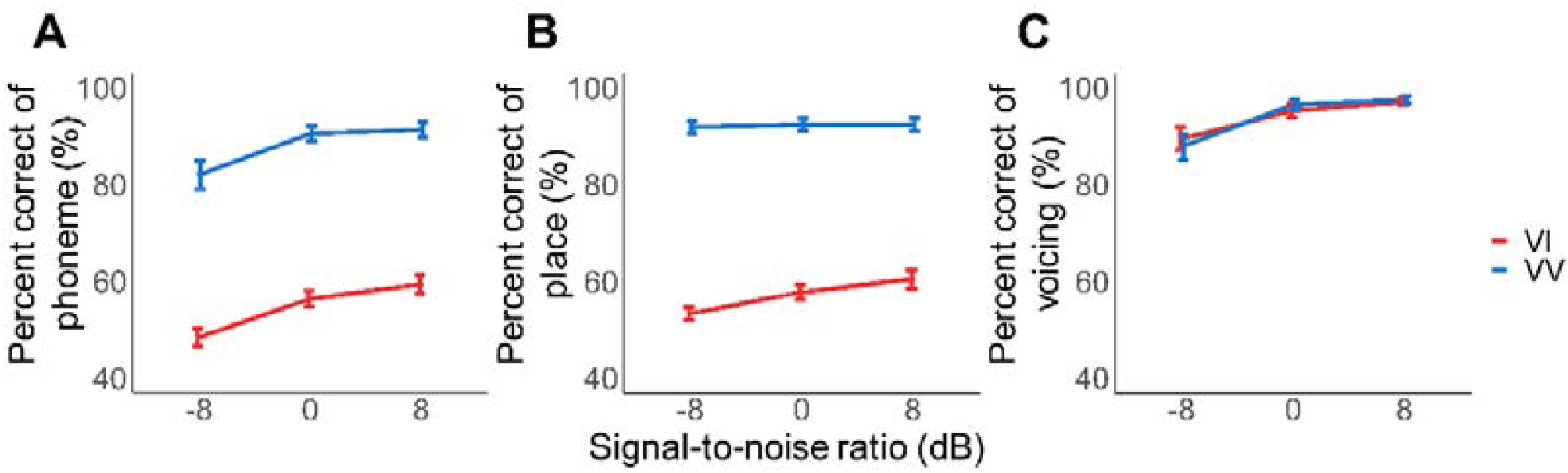
Behavioral performance. Mean percent of correct in identifying phonemes (A), the place of articulation feature (B) and the voicing feature (C) under visual valid (VV, blue line) and visual invalid (VI, red line) conditions.

Furthermore, a multiple linear regression analysis took both the visual enhancement (defined as VV - VI) of place recognition and that of voicing recognition into account in predicting the visual enhancement of phoneme recognition. Results showed that the visual enhancement of phoneme perception was remarkably explained by the visual improvement of place recognition (Supplementary Table 1, *β* = 0.89, *P* < 0.001) but was not related with the visual benefit of voicing recognition (Supplementary Table 1, *β* = 0.31, *P* = 0.249).

### Lip movements enhanced brain activity in sensory and motor areas

Univariate analyses showed significantly increased blood oxygen level-dependent (BOLD) activity in bilateral visual areas (left middle occipital gyrus (MOG), inferior occipital gyrus (IOG), middle temporal gyrus (MTG), fusiform gyrus (FFG), inferior temporal gyrus (ITG); right IOG, MTG and MOG), right STG and right prCG in the VV condition than in the VI condition (family-wise-error corrected *P* (*P*_*fwe*_) < 0.05, Fig. 2A and Supplementary Table 2), indicating stronger engagement of visual, auditory and motor regions by valid visual speech. In contrast, brain activity in bilateral lingual gyrus and left supplementary motor area (SMA) were weaker in the VV condition than in the VI condition. Consistent with prior findings (Du et al., 2014), SNR significantly modulated activity in auditory and speech motor areas, including bilateral STG, insula, triangular part of IFG, SMA, middle cingulate cortex, left MTG, MOG, AG, right SMG and postcentral gyrus (Fig. 2B and Supplementary Table 2). No significant interaction between visual validity and SNR was found (*P*_*fwe*_ > 0.05).

**Fig. 2.**
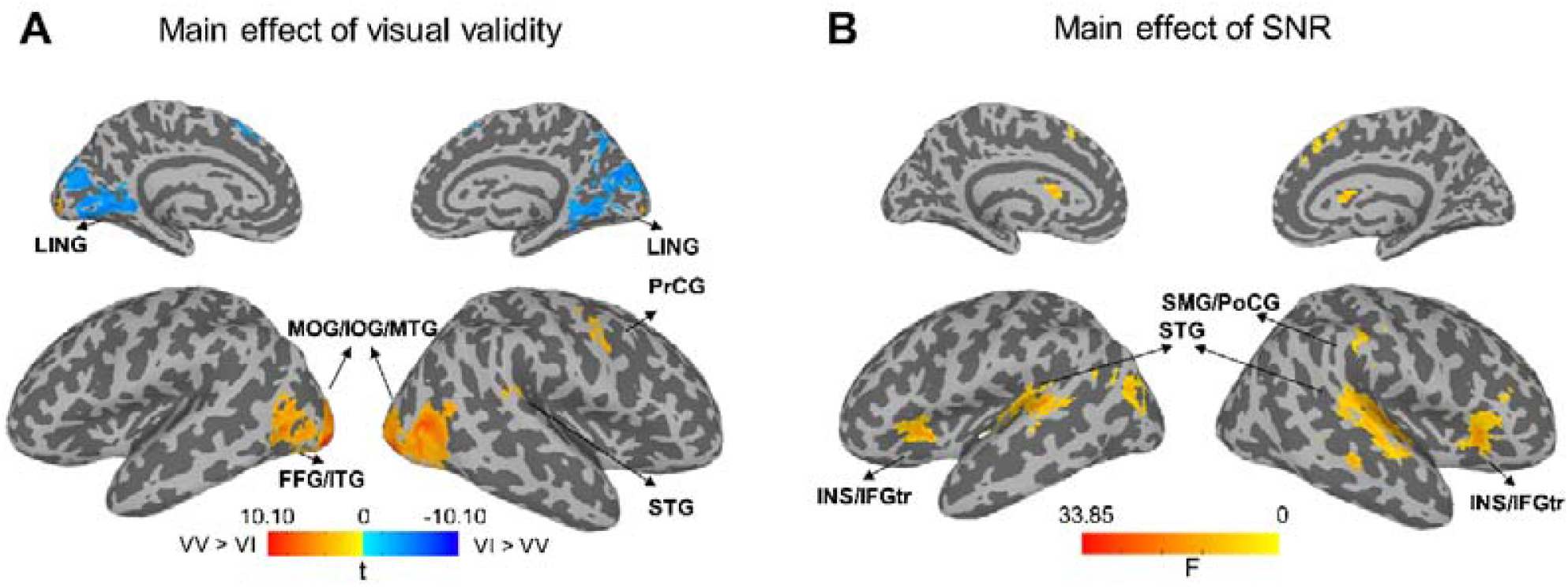
Main effects of visual validity and SNR on BOLD activity. (A) Regions where BOLD activity was modulated by visual validity (yellow: visual valid > visual invalid; blue: visual invalid > visual valid, *P*_*fwe*_ < 0.05). (B) Regions where BOLD activity was modulated by SNR (*P*_*fwe*_ < 0.05). FFG, fusiform gyrus; IFGtr, Inferior frontal gyrus, triangular part; INS, insula; IOG, inferior occipital gyrus; ITG, inferior temporal gyrus; LING, lingual gyrus; MOG, middle occipital gyrus; MTG, middle temporal gyrus; PoCG, postcentral gyrus; PrCG, precentral gyrus; SMG, supramarginal gyrus; STG, superior temporal gyrus.

### Lip movements sharpened the neural representations of speech phonemes and features

To examine the effect of visual validity on the neural representations of speech, we implemented MVPA in 50 individually defined anatomical regions of interest (ROIs) in both hemispheres (Fig. 3A) that were involved in audiovisual speech processing based on the previous review(Bernstein & Liebenthal, 2014). Support vector machine (SVM) classifiers were trained to decode the 4 phonemes on trial-wise fMRI response pattern ROI by ROI (see Materials and Methods).

**Fig. 3.**
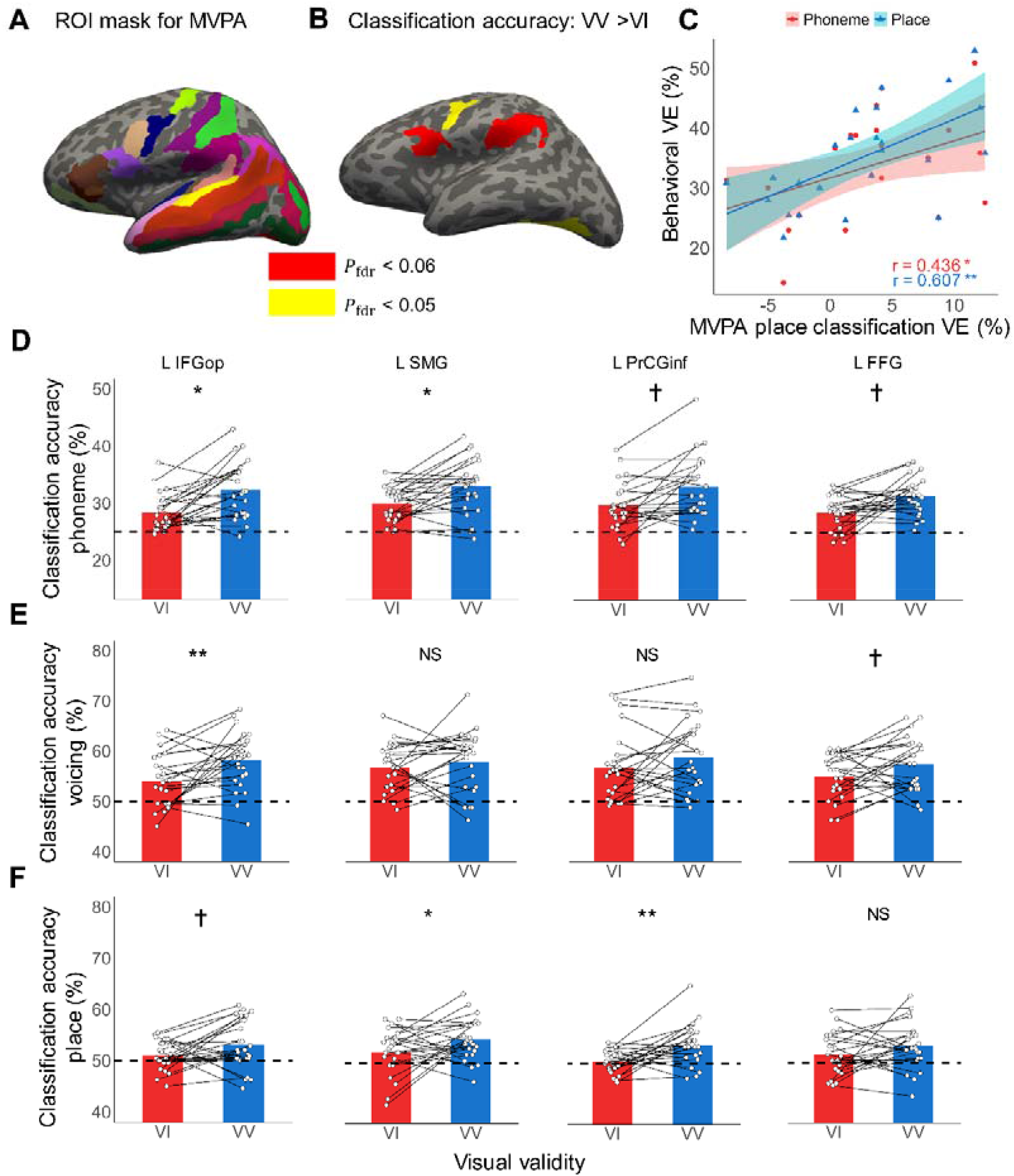
MVPA results. (A) Regions of interest (ROIs) in MVPA consisted of 25 left and 25 right anatomical ROIs implicated in audiovisual speech processing. (B) Regions where phoneme classification accuracy under the visual valid (VV) condition was higher than that under the visual invalid (VI) condition (red: *P*_*fdr*_ < 0.05; yellow: *P*_*fdr*_ < 0.06). (C) Correlation between visual enhancement (VE) of MVPA classification accuracy of place in the left SMG and visual enhancement of behavioral performance for recognition of phonemes (red) and place of articulation (blue), respectively. ** *P* < 0.01, * *P* < 0.05 by Pearson’s correlation. (D-F) The group mean and individual MVPA performance across SNRs in classifying phonemes (D), place of articulation (E) and voicing (F) in 4 ROIs. In panel D, * *P*_*fdr*_ < 0.05, † *P*_*fdr*_ < 0.06 by repeated-measures ANOVA with FDR correction. In panel E and F, ** *P* < 0.01, * *P* < 0.05, † *P* < 0.06, NS not significant by repeated-measures ANOVA without correction. Dash lines represent the chance level of classification. FFG, fusiform gyrus; IFG_op_, opercular part of inferior frontal gyrus; PrCG_inf_, inferior part of precentral gyrus; SMG, supramarginal gyrus.

A 2 (visual validity) × 3 (SNR) repeated-measures ANOVA found a significant improvement of classification accuracy of phonemes under the VV condition than the VI condition in the left IFG_op_ (*F*(1, 23) = 15.06, false-discovery-rate corrected *P* (*P*_*fdr*_) = 0.033, 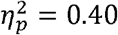) and the left SMG (*F*(1, 23) = 13.35, *P*_*fdr*_ = 0.033, 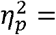 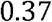) (red regions in Fig. 3B and 3D), although visual validity did not influence the overall BOLD activity in those regions. Additionally, a marginally significant improvement was found in the left PrCG_inf_ (*F*(1, 23) = 10.07, *P*_*fdr*_ = 0.053, 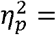 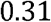) and the left FFG (*F*(1, 23) = 10.57, *P*_*fdr*_ = 0.053, 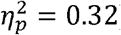) (yellow regions in Fig. 3B and 3D). The main effect of SNR and the interaction between SNR and visual validity were not significant in all ROIs (*P*_*fdr*_ > 0.6).

Since valid visual cues improved the recognition of place but not voicing at the behavioral level, we further investigated whether this was the case at the neural level. For those 4 ROIs showing significant or marginally significant visual benefit on phoneme classification, classification accuracy was recalculated according to the voicing or place of articulation feature as the same steps used in the behavioral analysis (see Materials and Methods).

Unprecedently, we observed diverse patterns of visual benefit on neural representations of articulatory-phonetic features in different regions. The left IFG_op_ showed a significant visual enhancement on representing voicing (*F*(1, 23) = 9.12, *P* = 0.006, 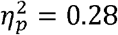) and a marginally significant visual benefit on representing place of articulation (*F*(1, 23) = 4.05, *P* = 0.056, 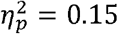). The left FFG only had a marginally significant visual enhancement on encoding voicing (*F*(1, 23) = 4.27, *P* = 0.050). In contrast, the left SMG and left PrCG_inf_ showed a significant visual enhancement on representing place of articulation (SMG: *F*(1, 23) = 4.54, *P* = 0.044, 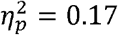; PrCG_inf_ : *F*(1, 23) = 11.51, *P* = 0.003, 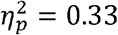) but an insignificant effect on encoding voicing (SMG: *F*(1, 23) = 0.67, *P* = 0.421, 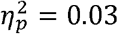; PrCG_inf_ : *F*(1, 23) = 2.00, *P* = 0.170, 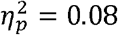) (Fig. 3E–F).

Next, we performed the correlation analysis to investigate the relationship between neural representations and behavior performance. We found that the visual enhancement of place classification in the left SMG was positively correlated with behavioral visual enhancement of place recognition (Pearson’s *r* = 0.61, *P* = 0.002), so as for phoneme identification (Pearson’s *r* = 0.44, *P* = 0.033) (Fig. 3C). No other correlation was found for any region (all Pearson’s |*r*| < 0.32, *P* > 0.125).

### Lip movements tightened the connection between dorsal stream areas and sensory cortices

We further conducted the DCM analysis to explore the effect of visual validity on the effective connectivity among audiovisual speech processing areas (Fig. 4A). Based on the univariate and MVPA results as well as the previous review(Bernstein & Liebenthal, 2014), the left SMG/AG and left speech motor areas (IFG_op_ and PrCG_inf_) were selected as amodal hub regions in the dorsal stream, and the left auditory cortex and visual cortex were included as sensory areas (see Materials and Methods).

**Fig. 4.**
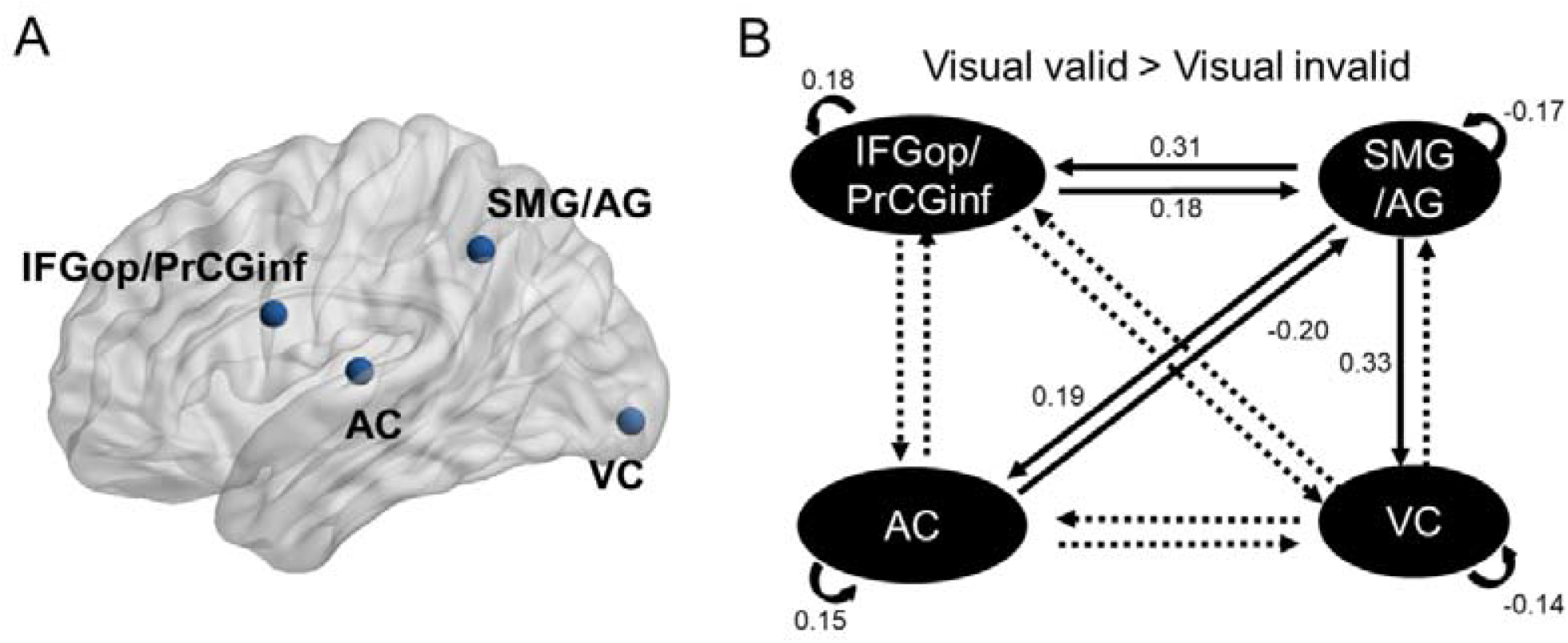
Dynamic causal modelling results. (A) Regions of interest in DCM. (B) The effective connectivities that were modulated by visual validity. Solid lines: *P*_*fdr*_< 0.05, dashed lines: *P*_*fdr*_> 0.05. Numbers represent averaged parameter estimates. AC, auditory cortex; AG, angular gyrus; IFG_op_, opercular part of inferior frontal gyrus; PrCG_inf_, inferior part of precentral gyrus; SMG, supramarginal gyrus; VC, visual cortex.

As shown in Fig. 4B and Supplementary Table 3, valid visual cues increased bidirectional connectivity between SMG/AG and speech motor areas (SMG/AG to speech motor areas: *t*(23) = 3.06, *P*_*fdr*_ = 0.015; speech motor areas to SMG/AG: *t*(23) = 2.43, *P*_*fdr*_ = 0.041) and top-down modulation from SMG/AG to auditory cortex (*t*(23) = 3.18, *P*_*fdr*_ = 0.013) and visual cortex (*t*(23) = 6.06, *P*_*fdr*_ < 0.001). However, bottom-up connectivity from auditory cortex to SMG/AG was inhibited by valid visual cues (*t*(23) = −3.46, *P*_*fdr*_ = 0.011). In addition, self-inhibition increased in auditory cortex (*t*(23) = 2.54, *P*_*fdr*_ = 0.037) and speech motor areas (*t*(23) = 3.91, *P*_*fdr*_ = 0.006), but decreased in visual cortex (*t*(23) = −3.37, *P*_*fdr*_ = 0.011) and SMG/AG (*t*(23) = −2.62, *P*_*fdr*_ = 0.035) when visual cues became valid. These results indicate that auditory cortex and speech motor areas became less sensitive to inputs from other regions while visual cortex and SMG/AG became more sensitive to inputs from other regions with valid visual cues. Correlation analysis found that self-inhibition in auditory cortex was negatively correlated with behavioral visual enhancement of phonemes (Pearson’s *r* = −0.54, *P* = 0.006) and behavioral visual enhancement of place recognition (Pearson’s *r* = −0.53, *P* = 0.008); self-inhibition in visual cortex was negatively correlated with behavioral visual enhancement of place recognition (Pearson’s *r* = −0.46, *P* = 0.025); and connectivity from speech motor areas to SMG/AG was negatively correlated with behavioral visual enhancement of voicing recognition (Pearson’s *r* = −0.41, *P* = 0.044). No other connectivity-behavior correlation was found (all Pearson’s |*r*| < 0.4, *P* > 0.052).

### Left AF microstructure predicted functional visual benefits

To further explore the structure-function relationship, we dissected 3 segments (long, anterior and posterior segments) of the left AF from the HARDI data. The mean fractional anisotropy (FA) from the diffusion tensor imaging (DTI) model and the mean neurite density index (NDI) and orientation dispersion index (ODI) from the NODDI model were calculated for each fiber bundle. The 3 AF segments connect ROIs in the MVPA results and DCM (see Materials and Methods), making it reasonable to implement correlation analyses between functional results and corresponding structural indexes.

As shown in Fig. 5, we found that the ODI of the long segment of AF (lAF, directly connecting IFG_op_ / PrCG_inf_ and posterior STG/MTG) was positively correlated with the visual enhancement of MVPA phoneme classification accuracy in the left IFG_op_ (Pearson’s *r* = 0.52, *P* = 0.010) and the visual enhancement of connectivity from speech motor areas (IFG_op_ / PrCG_inf_) to the auditory cortex (Pearson’s *r* = 0.41, *P* = 0.044). Meanwhile, the FA of the left lAF showed a negative correlation with the visual enhancement of MVPA phoneme classification accuracy in the left IFG_op_ (Pearson’s *r* = −0.43, *P* = 0.032) but no relationship with the visual enhancement of connectivity from speech motor areas to the auditory cortex (Pearson’s *r* = −0.31, *P* = 0.14). This result was consistent with the relationship between FA and ODI that FA decreases with a larger orientation variability. Additionally, the visual enhancement of phoneme classification accuracy in the left IFG_op_ was positively correlated with the visual enhancement of connectivity from speech motor areas to the auditory cortex (Pearson’s *r* = 0.45, *P* = 0.026). No other correlation between functional indices and structural indices of AF segments (all Pearson’s |*r*| < 0.23, *P* > 0.278), nor between behavioral performance and structural indices (all Pearson’s |*r*| < 0.29, *P* > 0.164) was found.

**Fig. 5.**
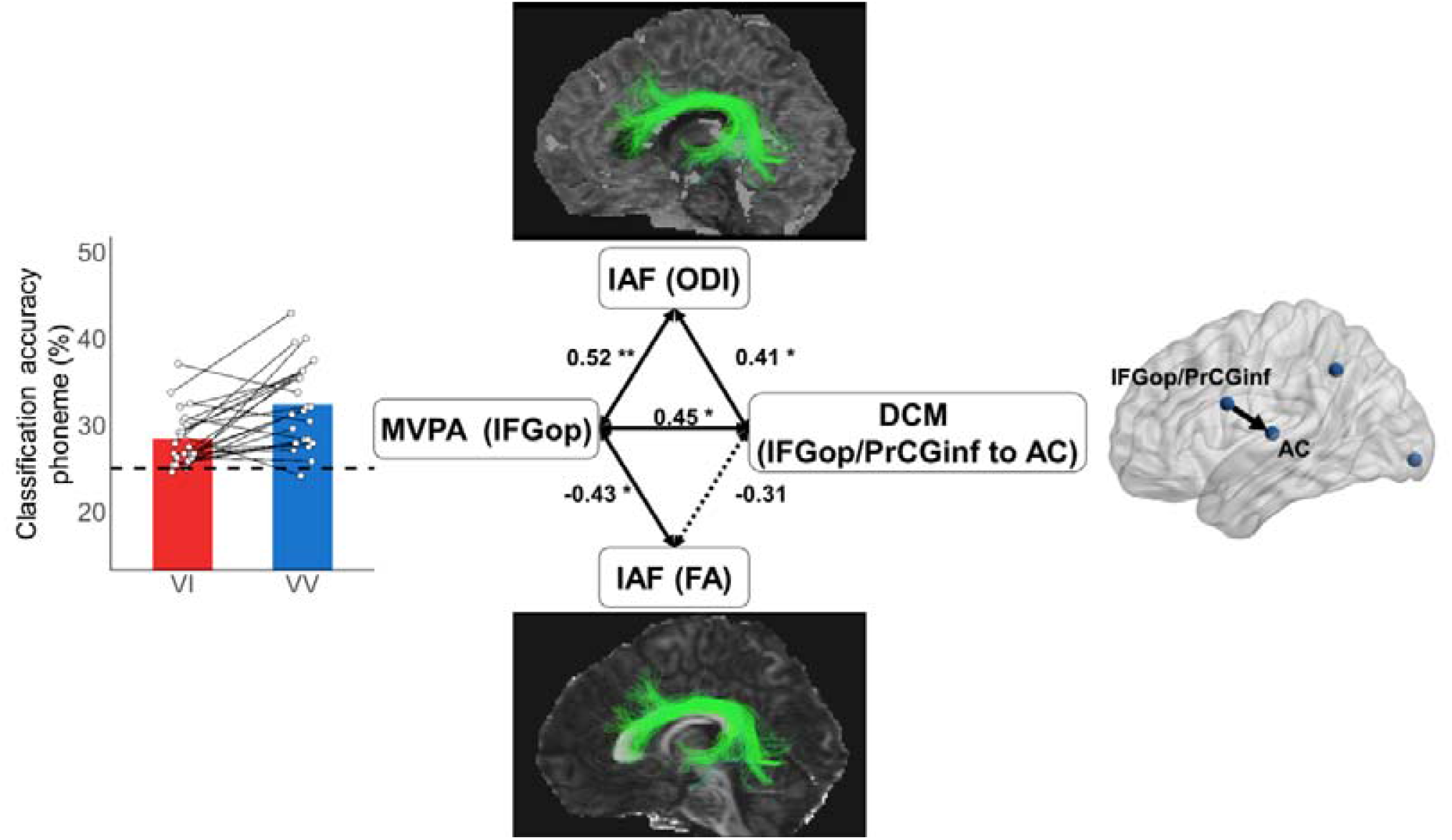
Correlations between the left AF microstructural properties and functional activity. The visual enhancement of phoneme classification accuracy in the left IFG_op_ correlated with the orientation dispersion index (ODI) and fractional anisotropy (FA) of the left long segment of arcuate fasciculus (lAF) and the visual enhancement of effective connectivity from speech motor areas to the auditory cortex. The visual enhancement of effective connectivity from speech motor areas to auditory cortex correlated with the ODI of the left lAF. Solid lines: *P* < 0.05, dashed lines: *P* > 0.05, ** *P* < 0.01, * *P* < 0.05 by Pearson’s correlation.

## Discussion

Visual lip movements improve speech-in-noise perception likely by constraining lexical interpretations and increasing the precision of prediction of the timing and content of the upcoming speech signal (Peelle & Sommers, 2015). Here we found that visual lip movements aided the behavioral recognition of the place of articulation rather than the voicing feature to promote phoneme perception in noisy conditions. MVPA results indicated that valid visual cues sharpened phoneme representations in speech motor areas (including the left IFG_op_ and PrCG_inf_) and the left SMG. Although it was not significant in the behavioral results, visual information enhanced the specificity of neural representations of voicing in the left IFG_op_, and improved the specificity of place representations in the left PrCG_inf_ and SMG. This fills the gap of knowledge that distinct dorsal stream regions are involved in processing different articulatory-phonetic features of audiovisual speech. In addition to regional neural representations, DCM analysis revealed significant visual modulations on the effective connectivity of the audiovisual speech network. With valid visual cues, SMG showed stronger bidirectional connectivity with speech motor areas and greater feedback connectivity to auditory and visual areas. Last but not least, the visual enhancement of phoneme specificity in the left IFG_op_ and the visual enhancement of effective connectivity from speech motor areas to the auditory cortex were correlated with each other, and both were correlated with the neurite architecture (orientation dispersion index) of the white matter tract connecting speech motor areas with auditory regions (the long segment of left AF). Our findings provide the first evidence that visual lip movements sharpen the neural representations of phonemes according to the place of articulation or voicing feature distinctively and promote the directed connectivity in the dorsal stream of speech processing, which is involved in sensorimotor integration (Hickok & Poeppel, 2007), and such visual benefits are mediated by neurite architecture of the dorsal stream fiber tract.

During speech processing, the speech motor areas (including Broca’s area and the left ventral premotor cortex) and the left pSTG/S are two candidate regions implicated in audiovisual integration (Peelle & Sommers, 2015). However, in our results, these two brain regions in the right hemisphere, rather than in the left hemisphere, showed stronger brain activation with valid visual cues. The stronger activation dose not equal to greater speech encoding ability, as only left speech motor areas showed greater specificity of speech representations with valid visual cues. This is consistent with previous findings that although both areas in the left could carry temporal information from auditory and visual modalities (Micheli et al., 2020), the left pSTG/S only represents redundant information of audiovisual speech, while the left motor areas represent synergistic information of audiovisual speech (Park et al., 2018). Similarly, MEG studies have found that both lip movements and speech envelope are tracked better only in left speech motor areas, but not in the left pSTG/S, with valid visual speech (Giordano et al., 2017; Park et al., 2016). Furthermore, a neuroimaging meta-analysis showed that the left pSTG/S is more steadily activated during conflicting audiovisual speech processing rather than validating speech processing (Erickson et al., 2014). Therefore, the left pSTG/S may be involved in solving the conflict between information from auditory and visual modalities, which was absent in the current study. In contrast, left speech motor areas may underlie the improved speech-in-noise perception with visual lip movements by enhancing neural representations of speech.

Consistent with previous findings (Grant & Walden, 1996), we found that lip movements significantly improved the identification of the place of articulation feature but not the voicing feature of speech. This supports the notion that the recognition of place and voicing is determined by the visual and auditory modality, respectively. Although we can easily discriminate voiced and voiceless consonants presented in noise merely by ear (90-100% correct, Fig. 1C), visual speech cues enhanced the neural encoding of voicing in Broca’s area (IFG_op_) without remarkable behavioral benefit. In contrast, the neural representations of place got improved by visual speech cues significantly in the left ventral premotor cortex (PrCG_inf_) and the left SMG, and marginally significant in the left IFG_op_. Although an audiovisual integration effect on encoding 19-dimensional phonetic features has been observed in a recent EEG study using continuous speech (O’Sullivan et al., 2021), to our knowledge, this is the first study that found a distinct visual enhancement effect of articulatory-phonetic feature representations in different brain regions. In the dual-stream model of speech perception, the dorsal stream can be further divided into the dorsal-dorsal stream that terminates in the premotor cortex (BA6, 8), and the dorsal-ventral stream that terminates in Broca’s area (IFG_op_, BA44) (Friederici, 2017; Rauschecker, 2018), implying the functional disassociation of the two speech motor areas. Speech production studies have found that the ventral premotor cortex represents articulatory gestures to a greater extent than phonemes, while Broca’s area represents both articulatory gestures and phonemes (Lotte et al., 2015; Mugler et al., 2018). It is posit that Broca’s area formulates the articulatory code which is passed to the premotor and motor cortices that subsequently implement the articulation during speech production (Basilakos, Smith, Fillmore, Fridriksson, & Fedorenko, 2018; Flinker et al., 2015; Long et al., 2016). In parallel, speech motor areas are hypothesized to generate articulatory predictions to compensate for degraded speech representations in the auditory cortex during speech perception when listening context (e.g., noisy, distorted speech) requires (Alain et al., 2018; Du et al., 2014, 2016; Nuttall et al., 2016; Pickering & Garrod, 2013; Skipper et al., 2017). Moreover, the left ventral premotor cortex exhibits articulator-specific engagement in speech perception (Liang & Du, 2018; Schomers & Pulvermüller, 2016). Under the audiovisual speech perception context, visual speech constrains the lexical competition by providing articulatory gestures, especially when the visual speech head start is processed before acoustic vocalization in most cases so that the articulatory prediction could be more precise (Karas et al., 2019; Peelle & Sommers, 2015). In particular, the left ventral premotor cortex is implicated in encoding the bottom-up lip movements (Ozker, Yoshor, & Beauchamp, 2018) besides receiving the top-down motor plans from Broca’s area, and extracting the synergistic feature of multimodal information (Park et al., 2018). Combining these findings, we hypothesized that Broca’s area and the left ventral premotor cortex might play a different role in audiovisual speech-in-noise perception. Specifically, Broca’s area might launch covert rehearsal and articulatory prediction to a greater extent and higher precision with visual speech cues, which would improve the neural differentiation of both voicing representations and place representations in Broca’s area. On the other hand, place of articulation rather than voicing is the major articulatory feature that visual lip movements provide to promote speech processing (Grant & Walden, 1996). Therefore, the left premotor cortex is assumed to automatically decipher the place of articulation information in lip movements by recruiting premotor subregions that control corresponding articulators, leading to enhanced topographical representations of place of articulation along the premotor strip. However, very few studies except Callen and his colleagues (Callan, Jones, & Callan, 2014) have investigated the distinct BOLD activity of the subregions in speech motor areas during audiovisual speech perception, further studies are needed to explore the detailed roles of speech motor subregions.

SMG is a multimodal brain region that anatomically and functionally connects auditory, visual and speech motor areas (Bernstein & Liebenthal, 2014; Binkofski, Klann, & Caspers, 2016; Donaldson, Rinehart, & Enticott, 2015). According to the dual-stream model of speech perception, the left SMG maps sensory representations into articulatory representations in speech processing (Gow, 2012; Hickok & Poeppel, 2007). Besides the speech processing network, SMG is also involved in the visual dorsal stream, which processes visuomotor sequences such as eye movements and hand movements (Basilakos et al., 2018; Meister, Wilson, Deblieck, Wu, & Iacoboni, 2007; Rauschecker, 2018). When speech is presented with lip movements, the left SMG would integrate visuomotor and acoustic information to promote the sensory-to-motor mapping. Therefore, the left SMG was recognized as a hub region in audiovisual speech perception where neural representations of phonemes and place of articulation were improved by lip movements. Notably, the visual enhancement of place representations in the left SMG predicted behavioral visual enhancement of phoneme recognition performance, which indicates that the representational changes in the left SMG may serve as a key neural substrate of the audiovisual benefit in speech processing.

The adding of visual speech cues not only sharpened speech representations in dorsal stream regions, but also strengthened the bidirectional effective connectivity of the dorsal speech stream (between SMG/AG and speech motor areas) and the top-down modulation from multimodal SMG/AG to unimodal sensory areas (auditory and visual cortices). The visual speech-induced stronger effective connectivity of the dorsal stream implies a greater extent of sensorimotor integration, which provides articulatory predictions to constrain phonological representations in sensory areas, to promote speech-in-noise perception (Du et al., 2014, 2016; Du & Zatorre, 2017; Hickok, Houde, & Rong, 2011; Hickok & Poeppel, 2007; Pickering & Garrod, 2013). An MEG study indeed found that the behavioral visual benefit is not predicted by changes in local speech entrainment but rather by enhanced effective connectivity between inferior frontal and temporal cortices (Giordano et al., 2017). In the current study, although we found no significant correlation between behavioral benefit and frontal-temporal connectivity, we revealed a positive correlation between the neural specificity of phonemes in the left IFG_op_ and the effective connectivity from speech motor areas to the auditory cortex. This result suggests that the better speech representations in frontal motor areas may lead to a stronger top-down constraint to auditory speech processing.

Another finding from the DCM analysis is that when a valid visual speech cue was presented, the auditory area became more self-inhibited (less sensitive to inputs from the network), but the visual area became less self-inhibited (more sensitive to inputs from other brain regions). This echoes a previous study (Nath & Beauchamp, 2011), in which the functional connectivity between the sensory cortex and the multisensory area is found reliability-weighted, that the multisensory region tends to be more strongly connected to the sensory area with more reliable information. That is, with valid visual cues, auditory information became less dominant to speech processing, and the visual cortex became more engaged in speech perception. Besides self-inhibition, the feedforward connectivity from the auditory area to the multimodal SMG/AG also reduced under the visual valid condition, supporting that auditory inputs became less weighted when visual cues are informative. Although we did not find a significant visual modulation effect regarding the connectivity from the visual cortex to other brain regions, weaker self-inhibition of the visual cortex correlated with stronger behavioral visual enhancement of place recognition, again demonstrating the increased contribution of visual modality.

Importantly, we further used in vivo NODDI technique to quantify the microcircuitry in terms of axon and dendrite complexity of the left AF, which is recognized as the neuroanatomic foundation of the dorsal stream in speech processing (Friederici, 2017; Hickok & Poeppel, 2007) and speech-in-noise perception (Li, Zatorre, & Du, 2021; Tremblay et al., 2019). The structure-function correlation analysis showed that the visual enhancement of effective connectivity from speech motor areas to the auditory cortex and the visual enhancement of phoneme representations in the left IFG_op_ were positively predicted by the ODI of the long segment of left AF, which directly connects the auditory cortex and speech motor areas. We also found a negative correlation between DTI-derived FA of the long segment of left AF and phoneme representations in the left IFG_op_. The opposite pattern between ODI and FA is consistent with our knowledge that the larger FA is correlated with greater NDI and lower ODI (Zhang et al., 2012). Note that, NODDI has been widely used in clinical populations, and previous studies have revealed that NODDI-derived ODI and NDI of white matter (Fu et al., 2020) and grey matter (Nazeri et al., 2015; Vogt et al., 2020) provide more specific microstructural indices than DTI-derived FA and macrostructural changes to cognitive aging, mild cognitive impairment and Alzheimer’s disease. The more robust structure-function correlation observed by ODI than by FA in the current study supports the above notion. However, NODDI has very recently been introduced to human cognitive neuroscience to investigate the relationship between brain morphometry and cognition in normal participants. One study has found a correlation between higher neurite density of the left planum temporale and higher temporal precision and shorter latency of auditory speech perception (Ocklenburg et al., 2018). In other two studies using HARDI data to estimate the fiber orientation distributions (FOD), the apparent fiber density (AFD) and the number of fiber orientations (NuFO) of the left AF are correlated with speech-in-noise perception criterion(Tremblay et al., 2019), and the AFD of the right AF is associated with EEG effective connectivity along the AF (Oestreich, Randeniya, & Garrido, 2019). To the best of our knowledge, this is the first study to introduce white matter neurite imaging to speech processing research and to investigate the relationship among neural representations, effective connectivity, and fiber neurite architecture during audiovisual speech perception. Although our analyses were rather exploratory, our findings imply that the higher dendritic complexity of the left AF may contribute to stronger benefits from the visual speech in enhancing neural specificity of phoneme representations and effective connectivity of the speech dorsal stream. This is the first evidence of the microstructural underpinning of functional performance in audiovisual speech-in-noise perception, and opens new avenue for future research.

Lastly, we did not find a significant interaction between visual validity and SNR on either BOLD activity or MVPA classification accuracy, which is unexpected since the visual benefit is assumed to be stronger in more noisy conditions than quieter conditions, i.e., the inverse effectiveness in audiovisual speech processing(Crosse et al., 2016). This may be caused partly by stringent correction procedure for multiple comparisons, and inappropriate SNR range to display the inverse effectiveness, as the performance even at the highest SNR (8 dB) in the visual invalid (auditory only) condition was relatively poor (~ 60% correct) inside the scanner.

In summary, we demonstrate that the speech dorsal stream is the key in visual enhancement of speech perception in noisy environments. Lip movements enhance both the specificity of phoneme representations and network connectivity of the dorsal stream to improve speech-in-noise perception. At the feature level, the visual enhancement on encoding place of articulation is revealed in the left ventral premotor cortex and multisensory SMG, while the visual enhancement on encoding voicing is observed in Broca’s area, providing novel evidence on interpreting finer roles of dorsal stream regions in articulatory-to-acoustic mapping during audiovisual speech processing. Importantly, this is the first report that the neurite orientation dispersion along the left AF can predict the visual benefits of neural representations and connectivity in the speech dorsal stream, pinpointing the microstructural property undergirding functional dynamics in multisensory speech processing. Our study paves the way for exploring local neural representations at different speech hierarchies, network dynamics, and microstructural characteristics underlying audiovisual speech perception.

## Materials and Methods

### Participants

Twenty-four young adults (19-28 years old, 12 females) participated in this study. All participants were healthy, right-handed, native Chinese speakers with no history of neurological disorder and normal hearing (average pure-tone threshold < 20 dB HL for 250 to 8,000 Hz) at both ears. All participants had signed the written consent approved by the Institute of Psychology, Chinese Academy of Sciences.

### Experimental design

The stimuli comprised 4 naturally pronounced consonant-vowel syllables (/ba/, /da/, /pa/, /ta/) uttered by a young Chinese female. The 4 syllables have 2 orthogonal articulatory features, voicing (voiced: /ba/ and /da/; unvoiced: /pa/ and /ta/) and place of articulation (bilabial: /ba/ and /pa/; lingua-dental: /da/ and /ta/). The utterances were videotaped by a Sony FDR-AX45 camera in a soundproof room. Then, they were digitized and edited on the computer to produce a 1-second video. Video digitizing was done at 29.97 frames/s in 1024 x 768 pixels. The pictures of the videos were cut, retaining the mouth and the neck part. The audio syllable stimuli were nearly 400ms in duration, low-pass filtered (4-kHz), and matched for average root-mean-square sound pressure level (SPL). The masker was a speech spectrum-shaped noise (4-kHz low-pass, 10-ms rise–decay envelope) that was representative of the spectrum of 113 different sentences by 50 Chinese young female speakers. The speech stimuli were presented at 90 dB SPL, and the SPL of the maskers was adjusted to produce different SNRs (−8, 0, and 8 dB). Audio stimuli were presented via MRI-compatible Sensimetrics S14 insert earphones (Sensimetrics Corporation) with Comply foam tips, which maximally attenuate scanner noise by 40 dB.

The experiment was a 3 (SNR: −8, 0 and 8 dB) x 2 (visual validity: valid and invalid) factor design. Matching lip movements videos and still lip pictures (the first frames of the matching videos) were presented with speech signals in the VV and VI conditions, respectively. In the fMRI scanner, subjects were instructed to listen to the speech signals, watch the mouth on the screen, and identify the syllables by pressing the corresponding button using their right-hand fingers (index to little fingers in response to /ba/, /da/, /pa/, and /ta/ in half of the subjects or the reverse order in the other half). Each subject completed 4 blocks of VI conditions and 4 blocks of VV conditions. The conditions were arranged in an ABBA or BAAB order, which was counterbalanced across participants. Each block contained 60 stimuli (20 trials x 3 SNRs), which were pseudo-randomly presented with an average inter-stimuli-interval of 5 s (4–6 s, 0.5 s step). Stimuli were presented via Psychtoolbox (Brainard, 1997).

### Behavioral analysis

We performed repeated-measures ANOVAs to investigate the effects of SNR and visual validity on phoneme-syllable identification or articulatory feature identification (voicing and place of articulation). Greenhouse–Geisser correction would be performed if the sphericity assumption was violated. Consistent with the previous study(Grant & Walden, 1996), if syllable /ba/ was recognized as /pa/, the response was correct for place and incorrect for voicing, while if syllable /ba/ was recognized as /da/, the response was correct for voicing, and incorrect for place. We further used a multiple regression analysis to determine the contributions of the visual enhancement on voicing and the visual enhancement on place to the visual enhancement on phoneme recognition. Visual enhancement was defined as the difference between the accuracy under the VV condition and the VI condition. Statistical analysis was conducted in R (R Core Team, 2017) with the package bruceR (Bao, 2020) and visualized using the package ggplot2 (Wickham, 2009).

### Functional imaging data acquisition and preprocessing

Functional MRI data were collected by a 3T MRI system (Siemens Magnetom Trio) with a 20-channel head coil. T1 weighted images were acquired using the magnetization-prepared rapid acquisition gradient echo (MPRAGE) sequence (TR = 2200 ms, TE = 3.49 ms, FOV = 256 mm, voxel size = 1×1×1 mm). T2 weighted images were acquired using the multiband-accelerated echo planar imaging (EPI) sequence (acceleration factor = 4, TR = 640 ms, TE = 30 ms, slices = 40, FOV = 192, voxel sizes = 3×3×3 mm).

The fMRI data were preprocessed using Analysis of Functional NeuroImages (AFNI) software (Cox, 1996). The first 8 volumes were removed for each block. For univariate analysis, the following preprocessing steps included slice timing, motion correction, aligning the functional image with anatomy, spatial normalization (MNI152 space), spatial smoothing with 6 mm FWHM isotropic Gaussian kernel, and scaling each voxel time series to have a mean of 100. The fMRI data were not spatially normalized, smoothed, and scaled for MVPA at the preprocessing steps.

### Univariate analysis

We conducted single-subject multiple-regression modeling using the AFNI program 3dDeconvolve. Six conditions of 4 syllables and 6 regressors corresponding to motion parameters were entered into the analysis. TRs were censored if the motion derivatives exceeded 0.3. For each SNR and visual validity, the four syllables were grouped and contrasted against the baseline.

We performed the group level analysis using the AFNI program 3dMVM. Two within-subject factors (visual validity, SNR) and their interaction were put into the model. Multiple comparisons were corrected using 3dClustSim (“fixed” version) with real smoothness of data estimated by 3dFWHMx (acf method) (Cox, Chen, Glen, Reynolds, & Taylor, 2017). 10000 Monte Carlo simulations were performed to get the cluster threshold (alpha□=□0.05 FWE corrected, uncorrected voxel-wise *P* < 0.005). Results were visualized onto an inflated cortical surface using SUMA with AFNI.

### ROI-based MVPA

We implemented MVPA in anatomically defined ROIs specific to each participant, thus no spatial normalization and smoothing was applied. We chose anatomical ROI-based MVPA rather than searchlight MVPA because we wished to preserve borders between spatially adjacent areas (e.g., IFG and STG) that were found to exhibit differential phoneme specificity at noisy conditions (Du et al., 2014, 2016). Freesurfer’s automatic anatomical parcellation (aparc2009 (Destrieux, Fischl, Dale, & Halgren, 2010)) algorithm was used to define a set of 148 cortical and subcortical ROIs from the individual’s anatomical image. We further divided STG into equational anterior and posterior portions, and divided prCG into equational dorsal and ventral parts. 25 ROIs in the left hemisphere that were closely related to audiovisual speech perception (Bernstein & Liebenthal, 2014) and the 25 counterparts in the right hemisphere were intersected with the Freesurfer mask to generate the 50 ROIs for MVPA. The classifiers were trained using SVM algorithm with a linear kernel. The cost parameter C was set to 1. The input feature was univariate trial-wise β coefficients that were estimated using AFNI program 3dLSS, which was recommended performing MVPA in fast event-related designs (Mumford, Turner, Ashby, & Poldrack, 2012). For each condition, the first level analysis of ROI-based MVPA was conducted within each anatomical ROI using the Decoding Toolbox (Hebart, Görgen, & Haynes, 2015). Twenty-fold cross-validation was used to evaluate classification performance, which was measured by the mean accuracy. Each fold contained a β coefficient of 1 trial of each syllable. We then conducted repeated-measures ANOVA with within-subject factors of visual validity and SNR in each ROI. Multiple comparisons were corrected with an FDR q = 0.05 using Benjamini–Hochberg procedure.

To further investigate the potentially different feature encoding visual benefits in different ROIs, for ROIs that showed significant or marginally significant visual enhancement on phoneme classification after FDR correction, we recalculated the classification accuracy according to the voicing and place feature with the same approach as the behavioral analysis. Repeated-measures ANOVA with within-subject factors of visual validity and SNR was performed to examine which feature representation was visually enhanced in each ROI.

### DCM analysis

We used DCM (Friston, Harrison, & Penny, 2003) analysis in SPM12 to assess effective connectivity among brain regions involved in audiovisual speech processing. Based on prior knowledge from the literature (Bernstein & Liebenthal, 2014) and our univariate and MVPA results, 4 ROIs (speech motor areas including IFG_op_ and PrCG_inf_, SMG and AG, auditory cortex and visual cortex) in the left hemisphere that were critical in audiovisual speech processing were selected in the DCM analysis. Although IFG_op_ and PrCG_inf_ showed different visual enhancement of feature representations in MVPA results, IFG_op_ and PrCG_inf_ were combined into one speech motor ROI in order to simplify the DCM model complexity. We identified the coordinates of each ROI according to the peak voxel of that region in the group-level activation under the VV condition. The group mean coordinates of ROIs were IFG_op_/PrCG_inf_ (−60, 6, 24), SMG/AG (−54, −52, 42), auditory cortex (−52, −18, 8) and visual cortex (−24, −94, −6).

We extracted the time series of each ROI according to the guideline (https://en.wikibooks.org/wiki/SPM/Timeseries_extraction). Since variation showing the maximum effect of interest between participants existed, we defined individual ROI as an 8 mm sphere centered on the individual peak activation voxel within a 15 mm sphere centered on the group peak voxel. This approach allowed individual ROIs to have slight variation between subjects, and be close to group peak coordinates. Voxels within the individual ROIs survived with the p < 0.05 uncorrected threshold were used to exclude the noisiest voxels within the ROIs. As suggested by the developers (Zeidman et al., 2019), if subjects with no voxel survived in an ROI existed, we increased the threshold with the step of 0.05 until all subjects got survived voxels in the ROI. Finally, we extracted the time series of survived voxels and used the first principal component of the extracted time series within the ROI in the subsequent DCM analysis.

We specified the modeling according to the DCM guide (Zeidman et al., 2019). DCM models the change of a neuronal signal x using the following bilinear state equation:

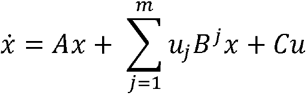

Matrix A denotes endogenous connectivity between modeled regions during baseline. Matrix B^j^ denotes the rate of change (in Hz) in connectivity between modelled regions with the j-th modulatory inputs. Matrix C represents how neuronal activity was influenced by the stimulus inputs. In the current study, the interested matrix was B^j^ that represents how effective connectivity among audiovisual speech processing regions was changed with valid visual cues in contrast to invalid cues. A positive parameter indicates that the connectivity increased. Conversely, a negative parameter indicates that the connectivity decreases. In addition, diagonal elements of the matrix, which indicate the intrinsic within-region self-inhibition, were also switched on. The more positive the self-connection parameter, the more inhibited the region, so the less it will respond to inputs from the network.

Since we assumed that valid visual speech would alter almost all the connections, we estimated a fully connected DCM for each subject using Bayesian model inversion. The main effect of the task (all trials) was set as the driving input to all ROIs (matrix C). The main effect of visual validity and SNR were set as modulatory inputs on the self-inhibition of each ROI (diagonal elements of matrix B^j^) and between-ROI connections (non-diagonal elements of matrix B^j^). Then, the spm_dcm_peb function was used to update the individual subject’ parameters using the group-level connection strengths as empirical priors, making summary statistics optimal. Finally, we extracted the expected connectivity parameters of matrix *B*^*j*^ from all participants. A one-sample t-test and FDR correction were performed to investigate the statistical significance of modulation (*P*_*fdr*_ < 0.05).

### Diffusion-weighted imaging data acquisition and preprocessing

Diffusion-weighted imaging (DWI) data were collected with following parameters: TR = 4000 ms, TE = 79 ms, voxel size = 1.5 × 1.5 × 1.5 mm, FOV = 192 mm, 64 gradient directions with two b values of 1000 s/mm^2^ and 2000 s/mm^2^, and 5 acquisitions without diffusion weighting (b = 0 s/mm^2^), which yielded the HARDI data.

The DWI data were pre-processed using MRtrix3 and FSL software (Jenkinson, Beckmann, Behrens, Woolrich, & Smith, 2012; Tournier et al., 2019). Preprocessing steps included denoising, unringing, eddy current and motion correction, and bias field correction using the N4 algorithm provided in Advanced Normalization Tools. Gradient directions were also corrected after eddy current and motion correction.

### HARDI tractography

Following the preprocessed step, tractography was conducted by MRtrix3 (Tournier et al., 2019). Firstly, 3-tissue (white matter, grey matter, and cerebrospinal fluid) response functions were obtained by the command “dwi2response dhollander” (Dhollander, Mito, Raffelt, & Connelly, 2019). Secondly, based on the response functions and preprocessed DWI data, we carried out multi-shell multi-tissue constrained spherical deconvolution (CSD) to estimate the FOD of each voxel. Thirdly, we performed the whole-brain tractography using second-order integration over FOD (IFOD2) probabilistic algorithm. Ten million streamlines were generated for each subject. Lastly, the command “tcksift” was used to filter the 10 million streamlines to 1 million streamlines (Smith, Tournier, Calamante, & Connelly, 2013).

AF has three segments: the long segment corresponds to the classical AF directly connecting the Broca’s area and the Wernicke’s area, the indirect anterior segment connects the Broca’s area and the Geschwind’s territory (the inferior parietal lobule) and the indirect posterior segment connects the Geschwind’s territory and the Wernicke’s area (Catani, Jones, & Ffytche, 2005). Three ROIs in the left brain (Broca’s area: IFG_op_ and PrCG_inf_ ; the Geschwind’s territory: SMG and AG; Wernicke’s area: posterior STG and MTG) were extracted from individual anatomical image parceled by Freesurfer, which were also used in MVPA. The extracted 3 ROIs were used to dissect the 3 segments of left AF according to the definition above in the native space.

### NODDI and DTI indexes calculation and correlation analysis

A NODDI model was fitted to each voxel of the preprocessed DWI data using AMICO python toolbox (Daducci et al., 2015) for each subject. The NODDI model is based on a 3-compartment tissue model (intra-cellular, extra-cellular and cerebrospinal fluid) and provides 3 indexes that are more specific to the white matter microstructure properties than FA index from the tensor model. NDI describes the number of neurites within a voxel, and ODI represents the variability of neurite orientations. For comparison, we also fitted a DTI model to each voxel of the preprocessed data using MRtrix3, generating an FA map for each subject. The relationship between FA and NODDI indexes is that FA increases with the increase of NDI or the decrease of ODI, and vice versa (Zhang et al., 2012).

We extracted the mean FA, NDI, and ODI along each AF segment for the correlation analysis. We performed the Shapiro-Wilk normality test to determine whether the variable was normally distributed. Then, we calculated Pearson’s correlation coefficients to assess the relationship between functional and structural results because all variables were normally distributed.

## Author contributions

L. Zhang acquired and analyzed the data. L. Zhang and Y. Du designed the experiment, interpreted the results and wrote the manuscript.

## Supplementary Materials

**Supplementary Table 1.** Results of the multiple linear regression analysis using the visual enhancement of recognition accuracy according to the place of articulation and voicing feature to predict the visual enhancement of phoneme identification accuracy.

|  | Visual enhancement (phoneme) |
| --- | --- |
| (Intercept) | 0.02 |
| Visual enhancement (place) | 0.89*** |
| Visual enhancement (voicing) | 0.31 |
| $R^2$ | 0.87 |
| Adj. $R^2$ | 0.85 |
| Num. obs. | 24 |

\* $p < .05$ , \*\* $p < .01$ , \*\*\* $p < .001$

**Supplementary Table 2.** Brain regions showing significant difference between visual valid and visual invalid conditions and a significant main effect of SNR (*P*_*fwe*_ < 0.05). AG, angular gyrus; Bi, bilateral; CALG, calcarine gyrus; CUN, cuneus; FFG, fusiform gyrus; HG, Heschl gyrus; IFGtr, Inferior frontal gyrus, triangular part; INS, insula; IOG, inferior occipital gyrus; ITG, inferior temporal gyrus; L, left; LING, lingual gyrus; MCC, middle cingulate cortex; MFG, middle frontal gyrus; MOG, middle occipital gyrus; MTG, middle temporal gyrus; PoCG, postcentral gyrus; PrCG, precentral gyrus; R, right; SFGmed, superior frontal gyrus, medial; SMA, supplementary motor area; SMG, supramarginal gyrus; STG, superior temporal gyrus.

| Brain Regions | Peak MNI coordinates |  |  | Peak t/F-value | No. of voxel |
| --- | --- | --- | --- | --- | --- |
|  | x | y | z |  |  |
| VV>VI |  |  |  |  |  |
| R IOG/MTG/MOG/FFG/ITG | 30 | -90 | -9 | 9.68 | 1046 |
| L MOG/IOG/MTG | -33 | -99 | -6 | 10.1 | 944 |
| L FFG/ITG | -42 | -45 | -21 | 6.64 | 132 |
| R PrCG/MFG | 54 | 3 | 57 | 4.77 | 59 |
| R STG | 78 | -45 | 18 | 4.66 | 40 |
| VV<VI |  |  |  |  |  |
| Bi LING/CALG/CUN | -9 | -54 | -3 | -6.55 | 1399 |
| Bi SMA/L SFGmed | 0 | 21 | 54 | -4.21 | 58 |
| SNR main effect |  |  |  |  |  |
| R STG/HG | 48 | -21 | 6 | 27.48 | 435 |
| L STG | -60 | -36 | 12 | 19.30 | 239 |
| L INS/IFGtr | -33 | 21 | 0 | 19.78 | 139 |
| R INS/IFGtr | 33 | 27 | 9 | 17.84 | 132 |
| L MTG/MOG/AG | -45 | -69 | 18 | 13.02 | 124 |
| L Caudate Nucleus | -6 | 6 | 15 | 33.85 | 114 |
| Bi SMA/SFGmed/MCC | 6 | 33 | 33 | 9.30 | 113 |
| R Putamen | 27 | -3 | -9 | 16.88 | 87 |
| L Putamen | -30 | -3 | 0 | 12.32 | 85 |
| R SMG/PoCG | 60 | -21 | 42 | 10.19 | 42 |

**Supplementary Table 3.** DCM results. Parameter estimates of the modulation effect of visual validity on the effective connectivity, the *t* and *P* values of one sample t-tests, and *P* values after FDR correction. AC, auditory cortex; AG, angular gyrus; IFG_op_, opercular part of inferior frontal gyrus; PrCG_inf_, inferior part of precentral gyrus; SMG, supramarginal gyrus; VC, visual cortex.

| | Estimate<br>(mean) | Estimate<br>(SEM) | $t$ | $P$ | $P_{\text{fdr}}$ |
| --- | --- | --- | --- | --- | --- |
| From AC to |  |  |  |  |  |
| AC | 0.149 | 0.059 | 2.54 | 0.019 | 0.037 |
| VC | 0.053 | 0.088 | 0.60 | 0.553 | 0.590 |
| SMG/AG | -0.198 | 0.057 | -3.46 | 0.002 | 0.011 |
| IFGop/PrCGinf | -0.107 | 0.056 | -1.92 | 0.068 | 0.109 |
| From VC to |  |  |  |  |  |
| AC | 0.142 | 0.087 | 1.63 | 0.117 | 0.170 |
| VC | -0.136 | 0.040 | -3.37 | 0.003 | 0.011 |
| SMG/AG | 0.006 | 0.065 | 0.09 | 0.926 | 0.926 |
| IFGop/PrCGinf | 0.176 | 0.132 | 1.33 | 0.196 | 0.241 |
| From SMG/AG to |  |  |  |  |  |
| AC | 0.187 | 0.059 | 3.18 | 0.004 | 0.013 |
| VC | 0.325 | 0.054 | 6.06 | <0.001 | <0.001 |
| SMG/AG | -0.172 | 0.066 | -2.62 | 0.016 | 0.035 |
| IFGop/PrCGinf | 0.312 | 0.102 | 3.06 | 0.006 | 0.015 |
| From IFGop/PrCGinf to |  |  |  |  |  |
| AC | -0.046 | 0.071 | -0.65 | 0.521 | 0.590 |
| VC | -0.087 | 0.062 | -1.41 | 0.171 | 0.228 |
| SMG/AG | 0.165 | 0.068 | 2.43 | 0.023 | 0.041 |
| IFGop/PrCGinf | 0.184 | 0.047 | 3.91 | 0.001 | 0.006 |

